# Learning sculpts microstructure in real time: evidence from dense temporal sampling of diffusion MRI

**DOI:** 10.64898/2026.06.24.734293

**Authors:** Deniz Kumral, Antonia Lenders, Monika Schönauer

**Affiliations:** Institute of Psychology, Neuropsychology, University of Freiburg, Freiburg im Breisgau, 79106, Germany; BrainLinks-BrainTools, University of Freiburg, Freiburg im Breisgau, 79106, Germany; Bernstein Center Freiburg, University of Freiburg, Freiburg im Breisgau, 79106, Germany

**Author notes:** D. Kumral, Institute of Psychology, Neuropsychology, University of Freiburg; Freiburg im Breisgau, Germany, Engelbergerstraße 41, Freiburg im Breisgau, 79106, Germany, **Email:**.

## Abstract

Learning induces rapid microstructural plasticity in the human brain, yet the precise temporal dynamics of these changes remain unclear. Using dense temporal sampling of diffusion-weighted MRI (DW-MRI) combined with task-based fMRI, we assessed microstructural changes throughout a declarative learning paradigm and subsequent rest. Seventy-four participants (36 females) learned image-location associations across four encoding-retrieval repetitions while undergoing interleaved functional and DW-MRI acquisitions. A matched control group (N=37, 21 females) underwent a similar imaging protocol without learning. Dense sampling of DW-MRI acquisitions (k=2146 across 22 time points in 127 min) revealed that learning-induced mean diffusivity (MD) decreases emerged shortly after learning onset and continued to develop during post-learning rest. The most robust and spatially consistent change was localized to the left middle occipital/temporal gyrus, a region also showing functional activation during encoding and retrieval. Linear mixed-effects modeling further confirmed a significant group-by-time interaction, with MD reductions in the left middle occipital/temporal gyrus emerging as early as ≈7 min after learning onset, becoming robust by ≈35-40 min, and persisting throughout the extended post-learning period, while controls showed no changes. Our findings demonstrate that learning-related microstructural plasticity unfolds continuously from encoding to offline consolidation, with learning-induced structural changes emerging in functionally engaged regions. Dense temporal sampling of DW-MRI offers a powerful approach to bridge functional activation and structural remodeling, providing evidence of *when* and *where* experience-dependent plasticity occurs during memory formation.

## Introduction

Learning induces not only transient functional changes in the brain, but also enduring structural modifications that support memory functions. Classic theories of systems consolidation propose that new experiences are first encoded by hippocampal–medial temporal circuits and then gradually integrated into distributed neocortical networks (McClelland et al., 1995; Frankland and Bontempi, 2005; Brodt and Gais, 2021). Traditionally, this process was thought to unfold slowly, requiring extended offline reactivation and reorganization (Klinzing et al., 2019). However, recent neuroimaging studies have indicated that microstructural remodeling can emerge within hours of a new experience (Sagi et al., 2012; Brodt et al., 2018; Tavor et al., 2020; Friedman et al., 2025), raising the question of *when the first* learning-induced microstructural changes can be observed in the human brain.

MRI has become central to assessing learning-induced changes at both the functional and structural levels (Villa et al., 2026). Functional MRI (fMRI), through the blood oxygenation level–dependent (BOLD) signal, has been used extensively to map the regions engaged in learning and memory (Brodt et al., 2016). However, while fMRI captures the hemodynamic correlates of neural activity, it does not reveal whether these regions undergo concurrent structural changes. Conversely, structural MRI (e.g., T1-weighted) has traditionally focused on long-term changes in gray matter volume (i.e., voxel-based morphometry) or cortical thickness accompanying extended training (Draganski et al., 2004; Zatorre et al., 2012; Hebscher et al., 2019; Kleinschroth et al., 2026). While these macroscopic changes are informative, they provide little insight into the microstructural processes occurring over shorter timescales.

Diffusion-weighted MRI (DW-MRI) provides an indirect measure of brain microstructure by assessing how freely water molecules move through tissue (Bihan, 1995, 2012). Mean diffusivity (MD), measuring the diffusive motion of water molecules, is particularly sensitive to microstructural changes (Assaf et al., 2019). Decreases in MD have been observed within hours of learning, including spatial navigation (Sagi et al., 2012; Tavor et al., 2013; Keller and Just, 2016), language learning (Hofstetter et al., 2017) and associative memory (Brodt et al., 2018), as well as motor learning (Jacobacci et al., 2020; Callow et al., 2023; Stee et al., 2023; Griffa et al., 2024; Friedman et al., 2025), and are thought to reflect processes such as synaptic changes, astrocyte functioning, and the expression of brain-derived neurotrophic factor (Johansen-Berg et al., 2012; Sagi et al., 2012), which is a marker for long-term potentiation. Although the aforementioned studies have demonstrated that learning induces microstructural changes in the human brain, the vast majority of research, including our own (Brodt et al., 2018), have used pre-post designs that cannot capture the temporal dynamics of plasticity (Villa et al., 2026). In contrast, continuous diffusion acquisition strategies, such as moving-average or sliding-window approaches, indicate that diffusion metrics can change within minutes (Darquié et al., 2001; Friedman et al., 2025). These findings suggest that learning-related plasticity is rapid and temporally heterogeneous across the brain. Nevertheless, it remains unclear whether these temporal dynamics extend to declarative memory. Specifically, do microstructural changes arise concurrently with learning, accumulate gradually across practice, or emerge only later, after learning has ended? How, *when*, and *where* are these changes stabilized in the brain?

In the present study, using dense temporal sampling of DW-MRI, we aimed to determine the spatiotemporal dynamics of learning-induced plasticity in humans, identifying *where* and *when* human brain microstructural plasticity arises. To investigate our aim, participants performed an image-location learning task which measures associative memory while undergoing an imaging protocol that interleaved traditional task-based fMRI runs with DW-MRI acquisitions. Following the task, participants underwent repeated DW-MRI acquisitions, allowing us to assess whether microstructural plasticity extended into the immediate post-learning phase. Crucially, the dense sampling of DW-MRI was designed to sensitively assess microstructural changes as they emerge and evolve throughout learning and post-learning rest phase. Rather than relying on pre-post comparisons, this design allowed us to determine when and where learning-induced microstructural changes emerge in functionally engaged regions.

## Materials and Methods

### Participants

Participants were recruited via the SONA online recruitment system, email, flyers, posters, and online advertisements. Potential participants were screened for eligibility through structured phone interviews. Eligible participants were between 18 and 35 years of age and were native German speakers or demonstrated at least B2-level proficiency in German. The exclusion criteria were current or previous treatment for psychological disorders, history of neurological conditions, untreated visual impairments, and contraindications for MRI. A total of 125 eligible participants were recruited, with 84 assigned to the learning condition and 41 to the control condition. Participants were excluded due to incidental findings (n = 2), falling asleep during data acquisition (n = 1), or withdrawing due to discomfort in MRI (n = 4; including 2 from the control group), resulting in an initial sample of 118 individuals: 79 in the learning group (*M* = 23.32 ± 2.96 years of age; 39 female) and 39 in the control group (*M* = 24.82 ± 3.50 years of age; 21 female). Additional exclusions were made due to poor image alignment (n = 3), excessive motion (mean framewise displacement averaged across sessions > 0.6 mm; n = 2), or incomplete datasets (n = 3). The final sample included 74 participants in the learning group (*M* = 23.22 ± 2.93 years; 36 females) and 37 in the control group (*M* = 24.70 ± 3.48 years; 21 females).

Participants were compensated at a rate of €10/hour for behavioral sessions and €15/hour for neuroimaging sessions, with total reimbursement ranging from €40 to €70, depending on the total time spent in the experiment. The study was conducted in accordance with the Declaration of Helsinki and approved by the local ethics committee (approval number: 21-1553) and all participants provided written informed consent. All tests were performed at the Department of Radiology, University Medical Center Freiburg, Germany.

### Experimental Procedure

Upon arrival, the participants received an MRI safety briefing and detailed instructions regarding the study procedure. They then completed a short tutorial of the image-location task and were randomly assigned to scanning group A, starting with fMRI acquisition during the first rehearsal block (N=37 in learning, N=18 in control), or B, starting with DW-MRI acquisition during the first rehearsal block (N=37 in learning, N=19 in control; see **Fig. 1**). The experimental session began with baseline acquisitions, including T1-weighted, T2-weighted, and DW-MRI (k=2) anatomical scans. In the learning group, after the baseline scans were measured, individuals completed N-back tasks, which serve as a localizer. This was followed by four repetitions of the image-location task, where task-based fMRI (k=2) and DW-MRI (k=8) were interleaved throughout the task (**Fig. 1**). After completing the image-location task, participants had a brief break before continuing the post-learning phase, which comprised resting-state fMRI (k=2) and DW-MRIs (k=12), followed by a final T1-weighted MRI. The final (fifth) retrieval session was conducted outside the MRI.

**Fig. 1.**
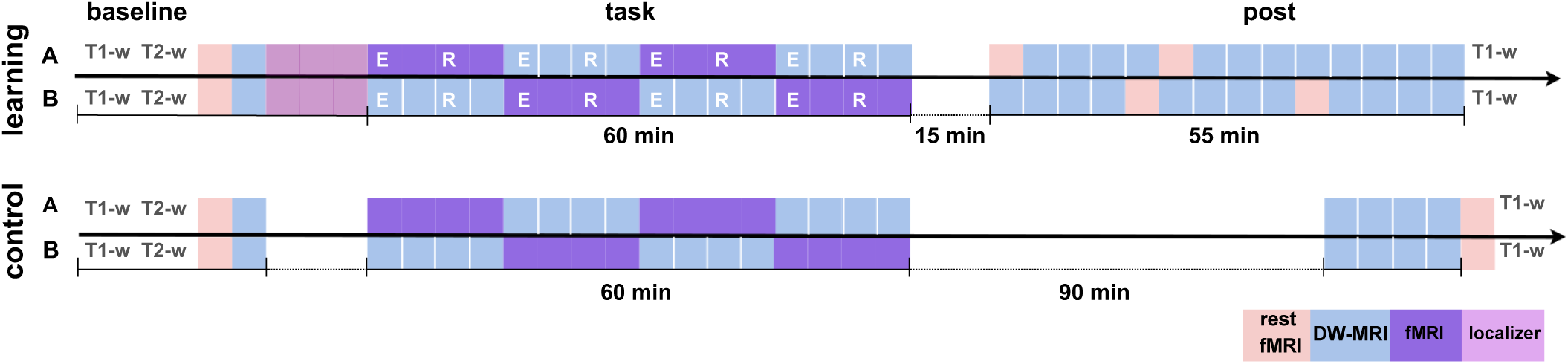
Experimental timeline and imaging protocol. Both scanning groups (A and B) underwent baseline MRI acquisitions, including T1-weighted (T1-w) and T2-weighted (T2-w) scans, followed by an image-location learning task consisting of alternating encoding (E) and retrieval (R) blocks (total duration: 60 min), with interleaved task-based fMRI and diffusion-weighted MRI (DW-MRI) acquisitions. After a short break (15 min), post-task resting-state fMRI and DW-MRI data were acquired (55 min), followed by a final T1-weighted scan. In total, 22 DW-MRI scans were acquired across baseline, task, and post-learning periods for more than 2 hours, enabling characterization of temporal dynamics of learning-induced microstructural changes in the brain. In the control condition, after identical baseline scans, participants remained at rest during the period corresponding to the learning task while interleaved resting-state fMRI and DW-MRI were acquired. Following a 90-min break, additional DW-MRI acquisitions and a final T1-w scan were measured. The control group completed 14 DW-MRI scans across comparable time windows.

The imaging procedure for the control group was largely equivalent to that of the experimental group. However, instead of performing the memory task, participants remained at rest during the phase when the task occurred for the learning group. Resting-state fMRI and DW-MRIs were acquired in an interleaved fashion following baseline imaging (k=2). After this phase, participants had a 90-min break and were instructed to avoid eating a full meal, studying, watching videos, exercising, or sleeping. Following the break, they returned to the MRI for the final session, which included four DW-MRI and T1-weighted MRI acquisitions.

### Image Location Task

During the experiment, participants were required to encode and recall item-location associations (Brodt et al., 2018). The experiment consisted of four encoding-retrieval repetitions and one additional post-scan behavioral retrieval session. During encoding, participants viewed 28 image pairs presented in random order. Each image was assigned a unique location on a 7×4 grid. The first image was shown full screen for 1.5 s to ensure that participants could perceive all visual details, then was moved onto its grid location in a continuous animation over 1 s, and remained visible there for 3 s. The second image (target) was then presented with the same durations; during the zoom-in period of the second image, the first image remained visible at its grid location, but was removed once the second image reached its position, such that the two images were not displayed together on the grid at the end of the trial (**Fig. 2**). The inter-trial intervals ranged from 1.7 to 2 s. Each image appeared in two different pairs, once as the first item and once as the second item. In the cued retrieval phase, the first image of each pair was presented for 2.5 s full screen before moving to its location on the grid. Participants navigated the grid using two button boxes (index and middle fingers of both hands for up, down, left, and right movements), with a red frame indicating the current position. Confirmation of the second image’s location was made by pressing a separate button with the pinky finger, turning the red frame white. The participants then used a red cursor to indicate the location of the second image. Confirmation was made by pressing a separate button. Each trial lasted up to 11.5 s, after which the cursor’s position was automatically recorded. All 28 pairs were tested once per retrieval session in a randomized order. At pseudorandom intervals, participants completed an active baseline task, classifying digits (1-9) as odd or even within a 2-sec response opportunity (Stark and Squire, 2001). Each baseline period lasted ~7.5 s, and digits were not repeated consecutively. The task was programmed using PsychoPy 2022.2.4 and was projected onto an MRI-compatible screen at the back of the scanner bore and viewed by participants via a mirror attached to the head coil. After scanning, the participants completed a fifth retrieval run on a desktop computer, again identifying the second-image locations.

**Fig. 2.**
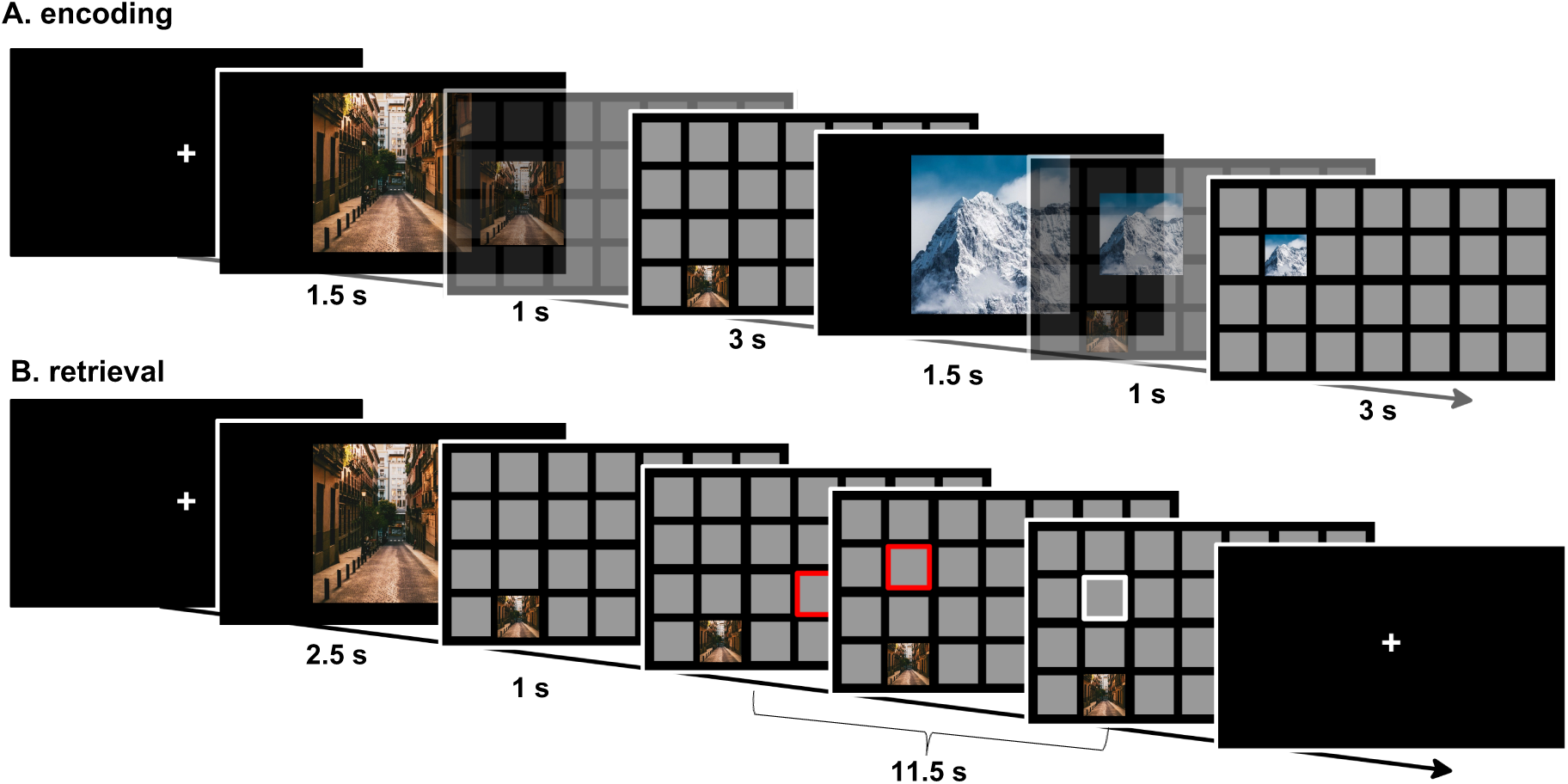
Schematic depiction of encoding and retrieval trials in the image-location task. Participants learned and recalled image-location associations across four encoding-retrieval repetitions. **A.** During encoding, 28 image pairs were presented in a random order, with each image assigned to a unique position on a 7×4 grid and shown once as the first item and once as the second item. **B.** During retrieval, the first image cued the recall of its pair and participants indicated the location of the associated image using a button-controlled cursor.

*MRI Acquisition.* Brain imaging was performed on a 3T PRISMA MRI scanner (Siemens Medical Systems, Erlangen, Germany) with a standard 64-channel head coil at the University of Freiburg, Medizinisch Klinik, Hugstetterstraße 55, 79106, Freiburg, Germany. To minimize offline warping, an auto-alignment scout was used to automatically position the measurement volumes consistently across baseline and post-learning scans.

Structural images were recorded with T1-weighted 3D high-resolution structural MRI brain scans (baseline and post-learning) using a GRAPPA sequence (acceleration factor 2) with the following parameters: TR = 2300 ms, TE = 4.18 ms, TI = 900 ms, slice thickness = 1.0 mm, bandwidth = 150Hz/Px, FA = 9°, FOV = 256 mm, and voxel size = 1.0 × 1.0 × 1.0 mm^3^, with 176 slices collected for a total collection time of 5:30 min. Likewise, we measured T2-w 3D high-resolution structural scans using a GRAPPA sequence (acceleration factor 2) with TR = 3200.00 ms, TE = 564.00ms, FA1 = 90°, FA2 = 120°, slice thickness = 0.8 mm, FOV = 256 mm, bandwidth 744Hz/Px, with a voxel size 0.8 × 0.8 × 0.8 mm^3^; 208 slices collected for a total collection time of 5:57 min.

The functional images were acquired via an echo-planar sequence (EPI) using the following parameters: TR = 1750 ms, TE = 30 ms, FA = 68°, FOV = 220 mm, imaging matrix = 88 × 88, with voxel size = 2.5 × 2.5 × 2.5 mm^3^, slice thickness = 2.5 mm, echo spacing = 0.52 ms, bandwidth = 2272 Hz/Px, 69 slices with an interleaved slice acquisition. A gradient echo field map with the sample geometry was used for distortion correction (TR = 400 ms, TE 1 = 4.92 ms, TE 2 = 7.38 ms, FOV = 205 mm, slice thickness = 3.2 mm). Imaging data were organized according to the Brain Imaging Data Structure (BIDS; Gorgolewski et al., 2016) to standardize data storage and allow for the application of automatic preprocessing pipelines.

DW-MRI scans were obtained with b-values 0 - 1000 s/mm^2^, in 30 whole-sphere gradient directions; 6 interspersed b0 images; TR = 2500.00 ms, TE = 53.00 ms, FOV = 220 mm, voxel size 2.5 × 2.5 × 2.5 mm^3^, partial Fourier ⅞, slice thickness 2.5 mm with 56 slices with a total time of 1:55 min. To enable use of FSL’s (Jenkinson et al., 2012) distortion correction tools topup and eddy, we acquired consecutively anterior-to-posterior and posterior-to-anterior phase encoding directions, totaling 44 DW-MR images (4 baseline, 16 learning, 24 post-learning) for learning and 28 DW-MR images (4 baseline, 16 learning rest, 8 post-learning) for control group.

### Structural and Functional MRI Data Preprocessing

Anatomical and fMRI data were converted to BIDS format (Gorgolewski et al., 2016) and later preprocessed using fMRIPrep 21.0.1 (Esteban et al., 2020) and Nipype (Gorgolewski et al., 2011). Briefly, the T1-weighted images were corrected for intensity non-uniformity, skull-stripped, and spatially normalized to MNI152NLin2009cAsym space using advanced normalization tools (ANTs; Avants et al., 2011). For functional data, we applied slice-timing correction and head-motion correction with six degrees of freedom. The BOLD data were co-registered to T1-weighted images, and nuisance regressors (e.g., framewise displacement [FD], DVARS, and CompCor) were computed. Finally, the BOLD time-series were spatially normalized to standard space using ANTs. Additional preprocessing was performed using Python code that included removal of 4 volumes of each image and also spatial smoothing with a 6 mm full-width at half-maximum (FWHM) Gaussian kernel.

### DW-MRI Data Preprocessing

Preprocessing of DW-MRI involved motion and eddy-current distortion correction using FSL’s *topup* and *eddy*. For each time point, all 36 b0 images across the two phase-encoding directions (1 AP, 1 PA) were used, resulting in 22 DW-MRIs in the learning and 14 DW-MRIs in the control group. Tensor calculation was performed using FSL’s *dtifit*, based on eddy-rotated b-vectors. Because fractional anisotropy (FA) images contain more structural landmarks, the registration was estimated on the FA data. Each participant’s FA image was co-registered to their high-resolution T1-weighted anatomical image (from *fMRIPrep*) using a 7 degrees-of-freedom linear registration (FSL *flirt*). This FA-to-T1 transform was then concatenated with the fMRIPrep-derived T1-to-MNI transform (MNI152NLin2009cAsym), and the combined transformation was applied to the MD images in a single resampling step using *antsApplyTransforms*. After registration, the MD images were smoothed using a 6 mm FWHM Gaussian kernel.

### Statistical Analyses

#### Memory Performance in the Image-Location Task

Behavioral performance was quantified using i) the percentage of correctly recalled locations of the paired images and ii) a normalized Manhattan distance, defined as the response-to-target distance (horizontal and vertical steps) divided by the start-to-target distance, such that values <1 reflect movement toward and values >1 movement away from the correct location. To assess whether participants improved their memory performance over learning repetitions, we applied a within-subject repeated-measures ANOVA. To compare performance at the fourth and fifth retrieval sessions against the first, we conducted paired-samples *t*-tests. All tests were two-tailed with an α level of 0.05. Due to incomplete data, one participant was excluded from both the behavioral and fMRI analyses.

#### Univariate fMRI Analyses

Functional data were analyzed using SPM12 (Wellcome Department of Cognitive Neurology, London, UK, https://www.fil.ion.ucl.ac.uk/spm). The preprocessed BOLD time series were high-pass filtered with a 128 s cutoff. A two-level general linear model (GLM) approach was used. First-level analyses were conducted using fixed effects at the individual level. The conditions of interest, encoding, recall, and baseline, were modeled for each repetition. The encoding time was defined as lasting from the onset of the first item of the image pair to the offset of the second item, while retrieval trials were specified as the duration from the onset of the first item to one volume before the participant made their first cursor movement, thus reducing motion-induced noise. If participants did not move the cursor, a time limit of 11.5 seconds was used as the offset of the retrieval event. The baseline time was defined as the onset and offset of the odd-even task. BOLD responses were modeled by convolving the event timings with the canonical hemodynamic response function. Six motion parameters were included as nuisance regressors.

We spatially smoothed the first-level contrast images, baseline-subtracted per condition, with an 8 mm FWHM Gaussian kernel before entering them into the second-level analysis. Full factorial models were computed with the factors of repetition and scanning groups. To preserve statistical power across runs, all trials, regardless of retrieval accuracy, were included. For second-level analyses, we used a linear contrast across runs (+1 +1 +1 +1) to assess the general activity during encoding and retrieval. Significant results were reported following family-wise error (FWE) correction for multiple comparisons (*p*_FWE_<.05, k=20).

#### DW-MRI Group-level Analyses

To assess group-level microstructural changes, flexible factorial models were used with time, experimental groups, scanning groups, and individuals as factors applied to whole-brain MD analyses in the MNI space. A total of 2146 images were included in the model. For the learning group, a linear decrease contrast was defined across all 22 acquired sessions using monotonically decreasing weights ranging from +11 to −11. For the control contrast, weights were constructed to temporally align with the corresponding time points in the learning group. This strategy ensured that, at each contrast, the full available sample in both groups contributed to the estimation while preserving temporal correspondence between groups despite unequal acquisition timings. Based on these group-specific vectors, a group × time interaction contrast was computed by directly contrasting the learning-related decrease with the corresponding timing-matched control condition (learning > control). Significant results were reported according to the FWE correction at the voxel level for multiple comparisons (*p*_FWE_<.05, k=20). To control for differences in the field of view, a whole-brain mask was created for each participant and session by excluding voxels deviating more than ±1 SD, and this mask was applied to the group-level analyses.

To further explore the temporal trajectories of learning-induced MD changes across the whole brain, we contrasted the MD change from baseline (t_1_) to each subsequent time point (t_2_-_22_) against the corresponding change in the control condition. This was implemented as an interaction contrast (i.e., [learning_t_n_ − learning_t_1_] - [control_t_n_ − control_t_1_]), resulting in 21 comparisons (p_uncorrected_<.001, k=20). For these contrasts, the corresponding control scans were defined by using temporally corresponding session(s). For time points of t_11_-t_19_, where no temporally matching control sessions were available, the final DW-MRI acquisition (t_10_) during the learning interval was used. By subtracting the control-related change from the learning-related change, these contrasts isolate MD changes specific to learning while accounting for nonspecific temporal effects.

To quantify the spatial consistency of learning-dependent MD changes over time, we performed a voxelwise overlap analysis across these 20 statistical maps, starting from learning onset (t_3_) for all subsequently sampled timepoints (t_3_-t_22_). Each map was binarized, and the resulting binary maps were summed voxel-wise to generate an overlap count image indicating the number of maps in which each voxel was present. An overlapping cluster was then derived by thresholding the overlap count at ≥10 maps (i.e., presence in at least 50% of all contrasts).

To determine whether the overlapping cluster (i.e., left middle occipital/temporal gyrus) coincided with functional activity during encoding and retrieval, we used joint inference analyses. First, the DW-MRI image was downsampled to the spatial resolution of the fMRI (2.5 mm). We then binarized the thresholded encoding and retrieval statistical maps (*p*_FWE_<.05, k=20) by retaining voxels with values greater than zero. A conjunction map was computed by taking the voxel-wise logical AND across the three binarized maps, identifying voxels that were jointly present in the encoding, retrieval, and left middle occipital/temporal gyrus.

### Linear Mixed-effects Model

To test for time-resolved MD changes between learning and control conditions, we next used linear mixed models using the *lme4* package (Bates et al., 2015) in R (version 4.3.1; R Core Team, 2023) with individuals as a random effect. We first extracted the MD values from the left middle occipital/temporal gyrus (k=17 voxels) across 22 DW-MRI scans acquired for each participant, capturing 38 distinct time points across the experimental protocol spanning ≈127 min, representing the real acquisition times rather than acquisition indices. These time points included baseline measurements (t_1_-t_2_, <20 min), learning task periods (t_3_-t_18_, 0-61 min), and an extended post-learning rest phase (t_19_-t_38_, ≈74-127 min), allowing for a comprehensive characterization of the temporal dynamics in the brain microstructure.

We used two models to enable targeted testing of the group-by-time interaction effect.

1. The null model included the main effects of time (as a factor with 38 levels representing different measurement time points across the baseline, task, and post-task periods) and group (learning vs. control), along with the scan group (A vs. B). A random intercept for participant ID was included to account for repeated measurements within individuals.
2. The full model was identical to the null model, except for the addition of the group-by-time interaction term, which represented the effect of primary interest (group × time).
3. The difference between the full and null models was tested using the anova function and setting the argument test to the chi-square (χ^2^) test. Statistical significance was set at α = 0.05.
4. In the case of significant interactions, the effects at the individual levels of predictors were analyzed post-hoc using Holm correction from the *emmeans* package (Lenth and Piaskowski, 2017). Note that we were only interested in examining MD changes from baseline within each group separately. For each group, we compared all time points (t_2-38_) to the baseline (t_1_) using the *contrast*() function with the “trt.vs.ctrl” method.

### Association between Memory Performance and Microstructural Plasticity

We performed two different correlation analyses to assess the correlation between MD and behavioral performance. A repeated-measure correlation (Bakdash and Marusich, 2017) was used to assess whether changes in MD across repetitions during learning were associated with improvements in memory retrieval during learning. Given the design with two scanning groups (Group A, *N* = 36: 1^st^ and 3^rd^ retrieval; Group B, *N* = 37: 2^nd^ and 4^th^ retrieval), analyses were performed at the two relevant time points per participant, and participant-specific mean MD values were extracted. We then averaged the MD values separately for each participant based on repetition, grouping runs 1 and 3 (A) and runs 2 and 4 (B), corresponding to their respective retrieval sessions. Repeated-measure correlations were then conducted using the *rmcorr* package in R (Bakdash and Marusich, 2017), which accounts for within-subject dependencies in the data and ensures robustness to potential violations of normality and homogeneity of variance. To further investigate brain-behavior relationships, Pearson’s correlation was used to test whether microstructural plasticity was related to memory gain. Microstructural change was defined as the percentage change in MD between the baseline (t_1_) and post-learning session (t_22_), and behavioral change as the improvement in recall (percentage and Manhattan Distance) between the first and fifth repetitions. All statistical analyses were performed using R (version 4.3.1; R Core Team, 2023).

## Results

### Behavioral Performance

The repeated-measures ANOVA revealed a significant main effect of repetition on memory performance (*F*(4, 353) = 45.54, *p* < .001, η2 = .37 for percentage and *F*(4, 353) = 35.75, *p* < .001, η2 = 0.33 for normalized Manhattan Distance, **Fig. 3B**). Post-hoc paired t-test comparisons showed that the percentage of correct answers significantly increased from the first (*M* = 15.11, SD = 10.94) to the fourth (*M* = 67.66, SD = 29.80) retrieval run (*t*(72) = −17.03, *p* < .001), and also from the first to the fifth retrieval (*M* = 63.83, SD = 30.2, *t*(70) = −15.51, *p* < 0.001; **Fig. 3A**). Likewise, we observed that the normalized Manhattan Distance decreased from the first (*M* = 1.15, SD = 2.81) to the fourth (*M* = 0.42, SD = 0.42) retrieval run (*t*(72) = 16.038, *p* < 0.001; **Fig. 3B**) and also from the first to the fifth retrieval (*M* = 0.473, SD = 0.429, *t*(70) = 15.025, *p* < 0.001). As expected, participants improved in recalling the items of an image-pair and their respective locations over learning repetitions.

**Fig. 3.**
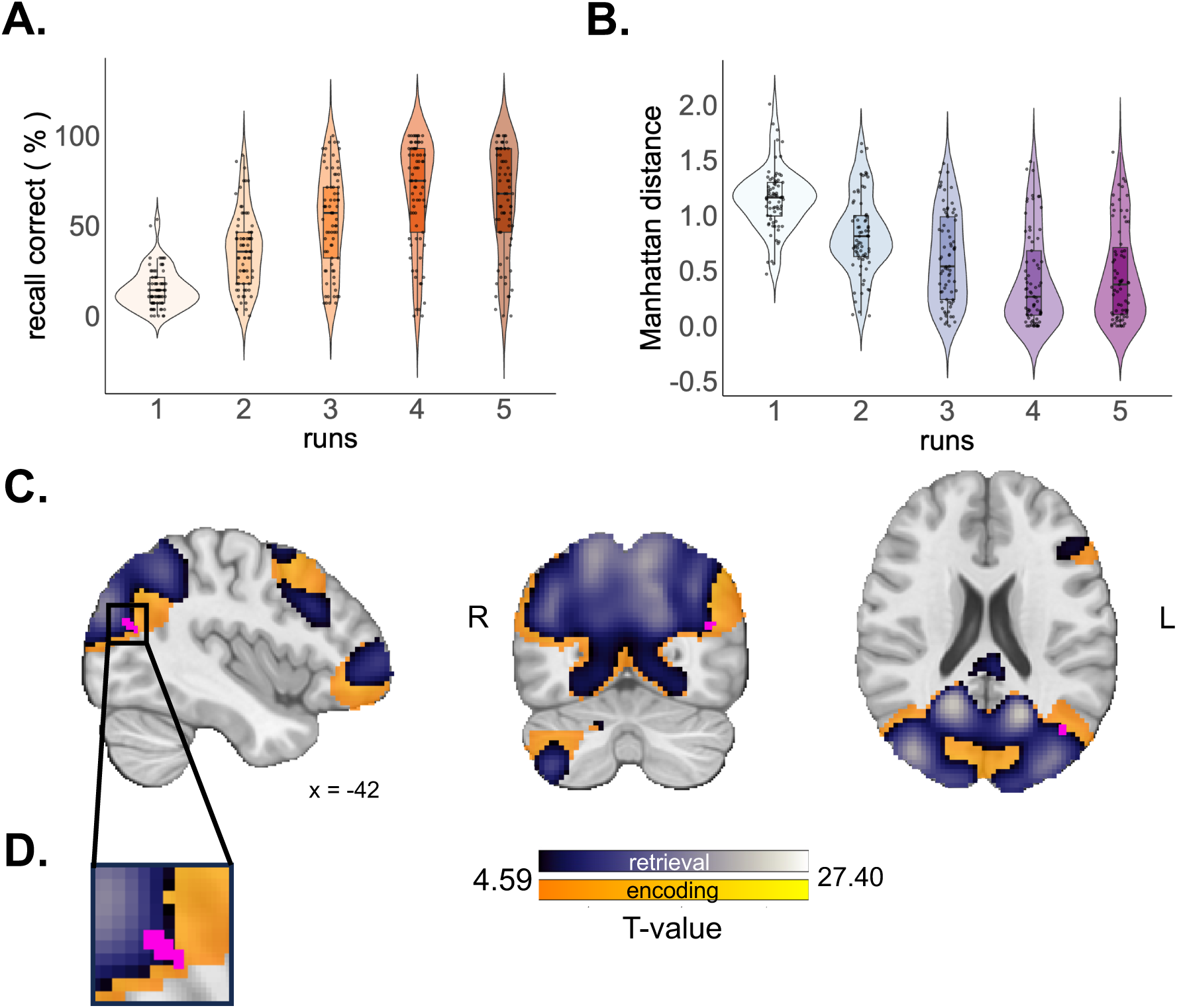
Behavioral performance across retrieval runs. Violin plots show the distribution of individual participant data for each run, while boxplots indicate the median and interquartile range, and dots represent individual participants. **A.** Percentage of correct responses increased across repetitions, while **B.** Manhattan Distance (normalized) decreased, indicating better recall accuracy. Repeated-measures ANOVAs revealed a significant effect of repetition on both measures (*p*<0.001). **C.** Statistical parametric maps illustrating general encoding-related activity (yellow–orange) and retrieval-related activity (blue–purple). **D**. Joint inference across task-based fMRI (encoding and retrieval) and the overlap map of Mean Diffusivity (MD) changes (each session [t_3_–t_22_] relative to baseline [t_1_]) revealed a shared cluster in the left middle occipital/temporal gyrus (dark pink; x = −41, y = −72, z = 21). Of the 17 voxels showing consistent MD change across time, 14 also overlapped with task-based functional activation, indicating that learning-related functional activity was accompanied by microstructural plasticity.

### fMRI Activity during Encoding and Retrieval

We conducted whole-brain analyses to identify brain regions associated with general functional activity changes during encoding and retrieval phases compared to an odd-even number judgement task. We observed widespread areas, including the occipital cortex extending to the temporal, parietal, and prefrontal cortices (*p*_FWE_<.05, k=20, **Fig. 3C**; Table S1–S2).

### Learning-related Microstructural Changes

We first defined a linear decrease contrast across all 22 acquired sessions using a whole-brain flexible factorial analysis with a group × time interaction, comparing the learning group against the control group. This revealed learning-specific decreases in MD (relative to control) in several brain regions, including the superior occipital gyrus, precuneus, cuneal cortex, superior parietal lobule, postcentral gyrus, cerebellum, middle occipital/temporal gyrus, and angular gyrus (**Fig. 4**, Table S3).

**Figure 4.**
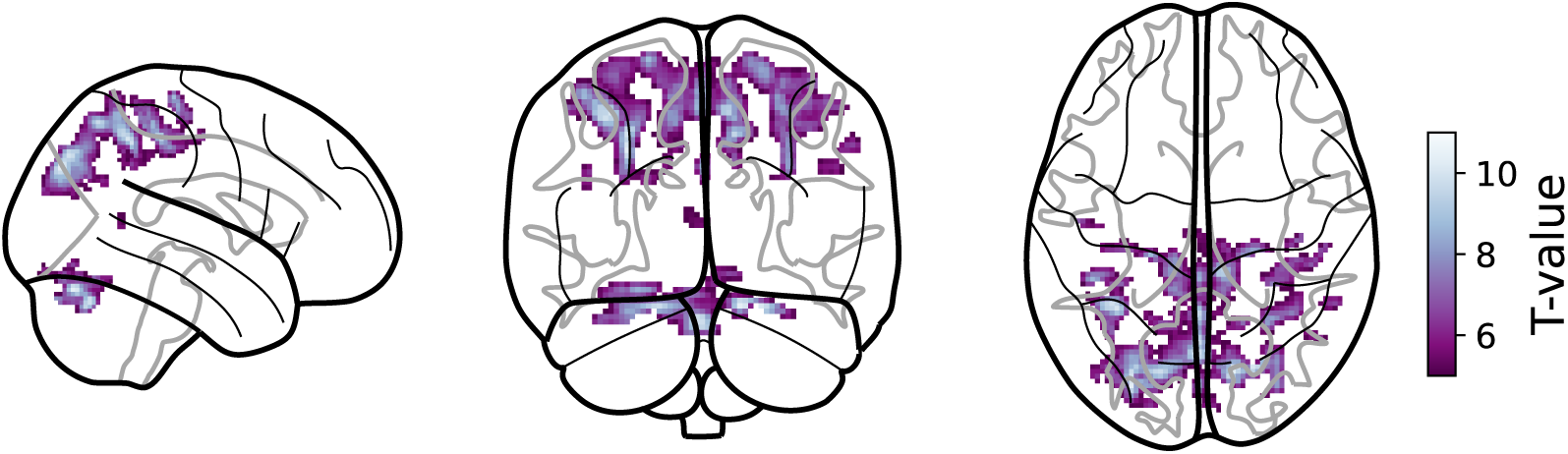
Learning-specific linear decreases in mean diffusivity (MD) across 22 sessions. We computed a group × time interaction (learning vs. control) using a whole-brain flexible factorial analysis with a linear decrease contrast. Relative to control, we found decreased MD in the learning group in several brain regions including the middle occipital-temporal gyrus, cerebellum, precuneus, superior parietal lobule, precentral gyrus, postcentral gyrus, cuneal cortex, angular gyrus, and lingual gyrus. Thresholded at *p* < 0.05, FWE-corrected with a minimum cluster size of k = 20 voxels.

We further assessed the temporal trajectories of learning-induced microstructural plasticity by contrasting the MD change from baseline (t_1_) to each subsequent time point (t_2_-t_22_) against the corresponding change in the control condition. As expected, at the baseline, no MD changes were detected (t_2_). During early learning, effects in the brain were sparse and mostly transient. From approximately the ninth session (t_9_), a cluster centred on the left middle occipital/temporal gyrus, falling within cluster identified in the overall linear-decrease analysis (**Fig. 4**, Table S3), emerged and persisted (**Fig. 5**). Beginning in the post-learning phase (t_11_), we observed MD decrease in several clusters, which increased steadily in both spatial extent and magnitude, peaking at the final acquisitions (t_19_-t_21_). These post-learning effects were predominantly posterior, encompassing the superior lateral occipital cortex, precuneus, and superior parietal lobule. Detailed statistics for the significant clusters of each contrast are provided in Table S4.

**Fig. 5.**
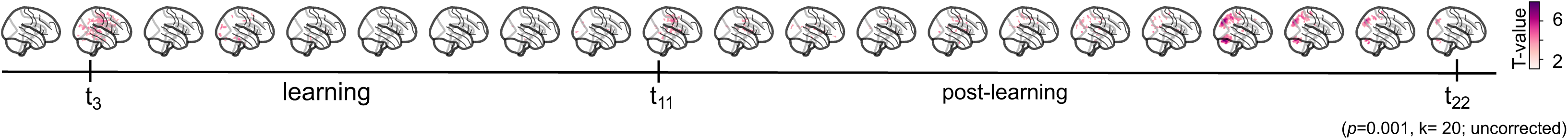
Temporal trajectories of learning-induced microstructural changes in the whole brain across sessions. We examined learning-induced changes in mean diffusivity (MD) at the whole-brain level by comparing baseline MD (t_1_) with each subsequent time point (t_2_–t_22_), relative to the control group, yielding 21 contrasts. Results revealed a progressive reduction in MD across several regions, including the precuneus, middle occipital/temporal gyrus, cerebellum, and lateral occipital cortex. Time points t_1_–t_2_ were defined as baseline, t_3_–t_10_ as the learning phase, and t_11_–t_22_ as the post-learning phase. All statistical parametric maps were thresholded at *p* < 0.001 (uncorrected) with a minimum cluster size of k = 20 voxels.

To evaluate the robustness of the observed effects, we assessed the spatial consistency of learning-dependent MD changes by overlaying the 20 thresholded contrast images spanning from learning onset to the end of the post-learning period (t_3_-t_22_) and computing their overlap, identifying voxels that showed MD reduction consistently across contrasts. Voxels surviving threshold in at least half of the maps (≥10/20) were retained, yielding a criterion that isolates regions reliably decreased across the time course. This analysis revealed a single robust cluster in the left middle occipital/temporal gyrus (k = 17; peak MNI coordinate [−41, −72, 21]). The cluster was consistently represented from the ninth (t_9_) to the twenty-first session (t_21_), indicating a spatially coherent and temporally stable locus of rapid MD reduction that emerged during the second half of the learning phase and persisted until the end of the measurement, 2 hours after learning onset (**Fig. 5**). The contribution of each contrast image to the overlapping cluster is summarized in Table S5. Joint inference analysis with fMRI further showed that the left middle occipital/temporal gyrus was activated during both encoding and retrieval (k=14, [x=−41, y=−72, z=21], **Figure 3D**).

### Fine-grained Temporal Dynamics of Microstructural Changes

To characterize how learning-induced microstructural changes in the left middle occipital/temporal gyrus unfolded over time, we applied linear mixed-effects models based on the real acquisition timings rather than acquisition indices. We observed a significant group-by-time interaction effect (χ^2^ = 59.33, df = 21, *p* < 0.001), indicating that the temporal trajectories of changes in brain microstructure differed significantly between the learning and control groups. The likelihood ratio test comparing the full model to the null model confirmed that the interaction term significantly improved the model fit (χ^2^ = 58.49, df = 21, *p* < 0.001; AIC full model = −44,823 vs. null model = −44,806). Type III Wald chi-square tests further revealed significant main effects of time (χ^2^ = 94.77, p < 0.001) and group (χ^2^ = 4.99, *p* = 0.025), while the scan group (A or B) showed no significant effect (χ^2^ = 0.02, *p* = 0.886).

To assess MD differences from baseline to the last session, we used post-hoc analyses, examining MD changes separately for each group (**Fig. 6**). While we observed no significant MD differences at any time in the control group (all *p* > 0.05, with effect sizes ranging from - 1.87 × 10^−6^ to 4.67 × 10^−6^), the learning group exhibited significant decreases in MD from baseline across the early task and post-learning rest periods. During the early task phase, the learning group showed small but significant decreases, beginning at ≈7 min (β = −4.67 × 10^−6^, *p* = 0.002) and continuing at 11.5 min (β = −3.41 × 10^−6^, *p* = 0.046), ≈20 min (β = −4.33 × 10^−6^, *p* = 0.005), ≈24 min (β = −3.52 × 10^−6^, *p* = 0.039), and ≈28 min (β = −3.41 × 10^−6^, p = 0.046).

**Fig. 6.**
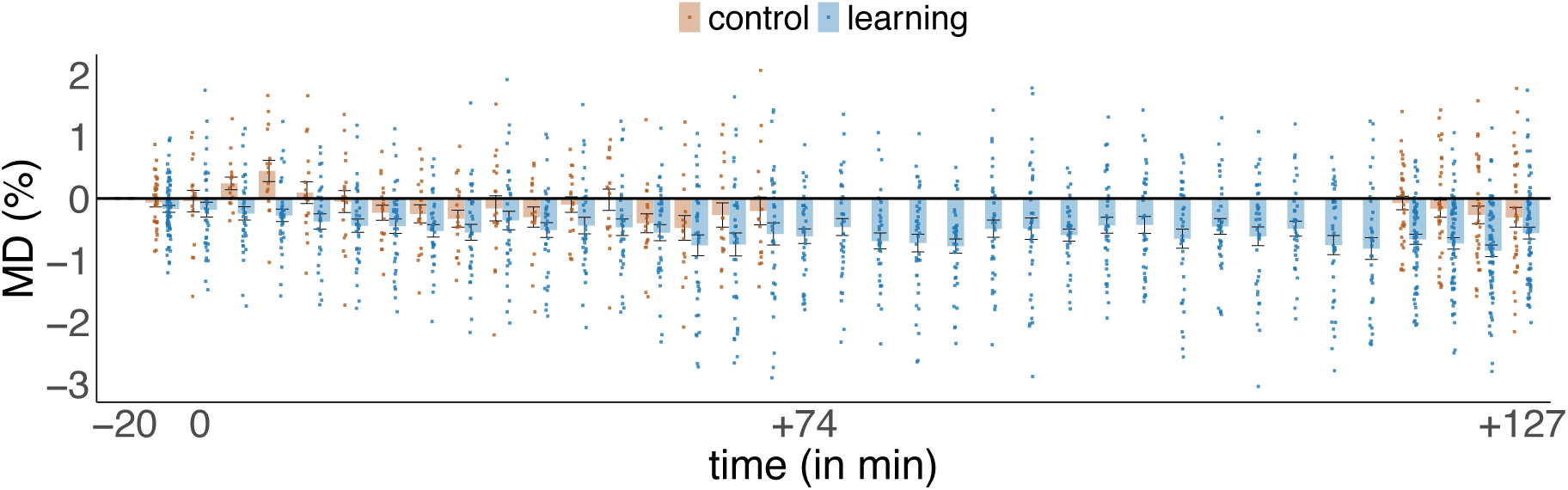
Fine-grained temporal dynamics of microstructural changes in left middle occipital/temporal gyrus. Mean Diffusivity (MD) changes from the baseline, expressed as a percentage. Each dot represents an MD change of each individual from their baseline (−20 min), while bar plots show the distribution of percentage changes at each time point. The post-hoc results from the linear mixed effect model indicate that the learning group demonstrates consistent MD decreases in the left middle occipital/temporal gyrus that emerge during early task phases and persist throughout the post-learning period, while the control group shows minimal deviation from baseline across all time points (see: Table S5). For visualization purposes, outliers (k=43 out of 2146) were removed using the interquartile range method, while all statistical analyses were performed on the complete data points (k=2146).

The strongest decreases during the task period occurred at a later time between ≈36-44 min (β ranging from −5.06 × 10^−6^ to −6.08 × 10^−6^, all *p* < 0.001) and ≈53-60 min (β ranging from −4.63 × 10^−6^ to −5.49 × 10^−6^, all *p* < 0.003). During the post-learning phase (≈74-127 min), all comparisons in the learning group showed significant decreases from the baseline (all *p* < 0.001). In summary, both the magnitude and the temporal persistence of MD decreases in the left middle occipital/temporal gyrus, alongside the lack of changes observed in the control group, provide strong evidence for rapid experience-dependent microstructural plasticity in the left middle occipital/temporal gyrus. This cluster also corresponds to the effects observed in the overall linear-decrease analysis (**Fig. 4**, Table S3). The results of the post-hoc pairwise comparisons and their corresponding times are presented in Table S5.

All results from whole-brain DW-MRI and fMRI statistical analyses can be found online in the public repository NeuroVault (https://neurovault.org/collections/24041/).

### Correlation between Memory Performance and MD

We first used repeated-measures correlation analyses to assess the relationship between MD changes in the left middle occipital/temporal gyrus and memory performance. We would expect stronger decreases in MD to coincide with increased memory performance if learning-induced microstructural plasticity was related to behavioral performance. When memory performance was quantified using the normalized Manhattan distance, the association was negative but did not reach statistical significance (r_rm_ = −0.22, *df* = 72, *p* = 0.060). An equivalent analysis using recall accuracy yielded similar results (r_rm_ = −0.21, *df* = 72, *p* = 0.063), suggesting decreased MD with increasing memory performance, although the effects remained at the trend level. Pearson’s correlation was used to examine the association between changes in learning-induced microstructural plasticity and memory retention (i. e., differences in memory performance between the first and fifth retrieval). No significant relationship was observed for either recall accuracy (r = −0.05, *p* = 0.63) or Manhattan distance (r = 0.10, *p* = 0.37).

## Discussion

In this study, we investigated the temporal dynamics of learning-induced microstructural plasticity in the human brain. Using a high-resolution DW-MRI protocol with 22 acquisitions over a 2-hour period, combined with task-based fMRI during an associative memory paradigm, we were able to assess *when* and *where* learning-induced microstructural changes occur in the human brain. We found that microstructural plasticity, indexed by a decrease in MD, emerged rapidly during learning, with reductions detectable within the first ≈7 min of the task. Importantly, these MD changes continued to develop and strengthen during the post-learning rest period, persisting for at least 2 hours thereafter. Notably, the most robust and spatially consistent effects were localized to the left middle occipital/temporal gyrus, co-localized with the functional activation we observed during both encoding and retrieval of the associative memory task.

A central finding of this study is that microstructural plasticity emerges rapidly in functionally relevant regions and persists into offline periods following the learning task. Learning-related MD reductions were observed across a broad set of regions including lateral occipital cortex, precuneus, and cerebellum, consistent with the distributed circuits typically engaged by visuospatial learning (Brodt et al., 2016, 2018). Remarkably, these MD reductions did not plateau at task offset; instead, they continued to expand and persist across the post-learning rest period, with the most robust effects localized to the left middle occipital/temporal gyrus. Linear mixed-effects modelling further revealed MD decreases emerging ≈7 min after learning onset and remaining detectable approximately 2 hours thereafter, consistent with prior reports demonstrating that MD changes can arise within minutes of learning (Sagi et al., 2012; Tavor et al., 2013; Hofstetter and Assaf, 2017; Brodt et al., 2018). Animal studies link these early changes to cellular mechanisms associated with long-term potentiation including astrocytic changes, increased BDNF expression, and elevated synaptic density (Blumenfeld-Katzir et al., 2011; Sagi et al., 2012; Garcia-Hernandez et al., 2022). Similarly, using continuous DW-MRI acquisition during learning, MD reductions have been reported to emerge within 10-30 minutes of training onset of a procedural learning and exhibit distinct temporal profiles across brain regions (Friedman et al., 2025). Extending this work, our findings demonstrate that associative learning similarly induces rapid microstructural changes that evolve rapidly in functionally relevant areas.

We further observed that MD decreases continued to strengthen during the post-learning rest phase, indicating that learning-related microstructural changes evolve beyond active task engagement. This trajectory aligns with the systems consolidation framework (Frankland and Bontempi, 2005; Tonegawa et al., 2018), wherein newly encoded memories undergo offline reprocessing and maturation over time, particularly in neocortical areas. Consistent with this, the emergence of further microstructural changes during rest suggests that offline periods support active neural and synaptic plasticity processes which strengthen and stabilize memory traces. Moreover, synaptic plasticity processes initiated during the task may continue to unfold, or emerge with a delay, after learning has ended. However, since microstructural changes were already detectable within ≈7 min of learning onset, such delayed processes cannot fully account for the trajectory, which instead spans both active learning and the subsequent rest phase. While the biological mechanisms underlying this offline plasticity remain unclear, they likely involve some combination of spontaneous neural replay, metabolic consolidation processes, glial responses to prior neural activity, and structural remodeling of synapses and dendrites. Understanding how these processes interact and contribute to the temporal dynamics of microstructural plasticity represents an important direction for future research.

A major aim of this project was to identify *where* in the brain learning-induced microstructural plasticity emerges most rapidly and is retained most robustly. The most consistent and spatially stable changes were evident in the left middle occipital/temporal gyrus. The middle occipital/temporal gyrus forms a key interface between visual perception and higher-order cognitive processing, transforming perceptual representations into increasingly abstract object-and meaning-based representations within the ventral visual stream and associated semantic networks (Patterson et al., 2007; Binder et al., 2009). It has been observed that occipito-temporal representations in visual working memory are shaped by feedback from parietal cortex (Xu, 2023), suggesting that short-term maintenance in this brain region might already reflect higher-order processing, potentially facilitating subsequent consolidation. It remains an open question whether such parietal-to-occipito-temporal interactions contribute to the longer-term consolidation observed here. Consistent with this account, our interactions analyses revealed MD decreases extending beyond the occipital/temporal cluster into higher-order parietal cortex, including the precuneus and adjacent posterior parietal regions. This offline parietal engagement aligns with prior evidence that the posterior parietal cortex rapidly forms enduring memory representations within hours of learning (Brodt et al., 2016, 2018; Kleinschroth et al., 2026) and that detailed item–context learning preferentially recruits posterior parietal areas (Klinkowski et al., 2025). Together, these findings are consistent with a circuit in which higher-order parietal regions interact with occipito-temporal cortex to support the stabilization of emerging category representations. The rapid plasticity detected in occipital/temporal gyrus thus may represent an early stage of more enduring structural remodeling. Finally, the spatial correspondence between functional activation and microstructural plasticity in this region suggests that learning-engaged regions undergo structural remodeling to store memory traces directly within the circuits that process the corresponding sensory input, consistent with the direct neocortical encoding of category information in early visual cortex and of episodic item–context information in posterior parietal cortex, reported recently (Klinkowski et al., 2025). Our findings thus challenge traditional models that assume memory representations must be abstracted to higher-order association areas, instead supporting a distributed organization where memories form in a more widespread manner along specialized processing streams.

Although the control group completed very similar DW-MRI acquisitions, enabling assessment of MD changes during comparable time windows, they did not undergo the same imaging acquisition procedure as the experimental group. In particular, the learning group performed a localizer task immediately before the learning phase, which may have contributed to a cluster of surprisingly rapid MD changes observed only briefly at the very onset of learning. These early changes could reflect transient effects associated with visual stimulation, potentially linked to BOLD activity (Darquié et al., 2001; Spencer et al., 2025). Moreover, the control group did not undergo all DW-MRI measurements in the post-task period, because we considered the strain of a 3-hour wakeful task-free rest period in the MRI scanner to be hardly tolerable. Consequently, this difference limits our ability to fully characterize time-dependent MD changes that may occur independently of learning, such as transient changes in water diffusion associated with neuronal activity (Darquié et al., 2001; Tsurugizawa et al., 2013) or non-specific effects like scanner drift (Vos et al., 2017). Therefore, the apparent strengthening of MD decreases across the post-learning period should be interpreted with caution. However, the fact that the learning group showed progressive, spatially specific MD decreases in the left middle occipital/temporal gyrus already during the learning phase, decreases that were absent in the control group during comparable time windows and that persisted beyond active learning, provides reassurance that our findings reflect genuine learning-related microstructural plasticity rather than non-specific time-dependent, measurement-dependent, or functional-activation-related confounds. Finally, the temporal resolution of DW-MRI is coarse compared to the sub-second timescale of synaptic plasticity processes. Nevertheless, our approach represents a substantial improvement over traditional pre-post designs. While Friedman et al. (2025) used a sliding-window approach to extract MD changes from continuously acquired DW-MRI during motor learning, our discrete acquisition strategy allowed us to synchronize 22 DW-MRI samples with specific task and rest phases, optimizing temporal sensitivity to state-dependent microstructural changes. Finally, fMRI and DW-MRI were acquired in an interleaved fashion rather than simultaneously, introducing temporal lags between measurements. While this prevented a continuous assessment of MD changes within participants, this design was necessary to capture both functional and structural progression across the learning period. The clear spatial overlap between functionally active regions and sites of microstructural plasticity, suggests that this limitation did not substantially impact our results. It is important to note that while MD is a well-established marker of tissue microstructure, it reflects a mixture of intra- and extracellular water compartments and does not isolate specific microstructural features such as dendritic complexity, synaptic density, or glial proliferation (Villa et al., 2026). Combining ultra-high-gradient DW-MRI with SANDI, recent work has dissociated a transient, task-wide expansion of cell bodies from a sustained, region-specific increase in cell-process density during motor learning, demonstrating how compartment-specific models can separate plastic from non-plastic processes (Griffa et al., 2026). Advanced DW-MRI models may therefore help identify which specific microstructural features are most affected by learning and further change across the subsequent offline consolidation phase.

In conclusion, using dense temporal sampling of DW-MRI, we demonstrate that learning-induced microstructural plasticity emerges rapidly within minutes of task onset and continues to develop during post-learning rest. The convergence of spatially consistent, temporally persistent microstructural plasticity and functional activity in the left middle occipital/temporal gyrus highlights its central role in associative memory. Our approach of densely sampling DW-MRI throughout learning and the post-learning rest phase, rather than simple pre-post comparisons, thus reveals the dynamic temporal unfolding of plasticity: initial emergence during early learning phases, strengthening during continued practice, and persistence through extended offline consolidation periods.

## Acknowledgments

We would like to express our gratitude to Tobias Debor, who helped with the experimental setup. We also thank all research assistants for helping with data collection.

This project was supported by Deutsche Forschungsgemeinschaft (DFG, German Research Foundation) – Project number: 426865207

## Supplemental Material

**Table S1.** Functional memory-related activity during encoding (*p*_FWE_<.05, k=20).

| Cluster size | T-value | x | y | z | Hemisphere | Region |
| --- | --- | --- | --- | --- | --- | --- |
| 17870 | 27.14 | 11 | -90 | 2 | Right | Occipital Pole |
| 3398 | 16.44 | -34 | 10 | 59 | Left | Middle Frontal Gyrus |
| 418 | 7.92 | -59 | -55 | -6 | Left | Middle Temporal Gyrus,<br>temporooccipital part |
| 385 | 9.68 | 28 | 12 | 52 | Right | Middle Frontal Gyrus |
| 337 | 13.02 | 11 | -50 | -48 | Right | Cerebellum |
| 293 | 6.78 | -19 | 48 | -18 | Left | Frontal Pole / Orbitofrontal cortex |
| 89 | 6.31 | 31 | 58 | 4 | Right | Frontal Pole / Middle Frontal Gyrus |

**Table S2.** Functional memory-related activity during retrieval (*p*_FWE_<.05, k=20).

| **Cluster size** | **T-value** | **x** | **y** | **z** | **Hemisphere** | **Region** |
| --- | --- | --- | --- | --- | --- | --- |
| 15521 | 27.22 | 14 | -92 | 2 | Right | Occipital Pole |
| 1983 | 21.34 | -24 | 2 | 59 | Left | Middle/Superior Frontal Gyrus |
| 778 | 15.92 | 24 | 2 | 56 | Right | Middle/Superior Frontal Gyrus |
| 587 | 15.08 | 14 | -50 | -48 | Right | Cerebellum |
| 317 | 11.81 | 38 | -70 | -51 | Right | Cerebellum |
| 765 | 10.92 | -42 | 55 | 2 | Left | Frontal Pole |
| 63 | 8.59 | -29 | 25 | -1 | Left | Insula / Orbitofrontal Cortex |
| 122 | 7.23 | -59 | -60 | -6 | Left | Fusiform Cortex / Lingual Gyrus |
| 102 | 6.64 | -12 | 10 | 9 | Left | Caudate |
| 26 | 6.02 | -6 | -88 | -36 | Left | Cerebellum |
| 26 | 5.48 | 8 | 12 | 9 | Right | Caudate |

**Table S3.** Learning-induced microstructural changes based significant group × time interaction effects: linear MD decrease in learning vs. linear MD increase in control group (*p*_FWE_<.05, k=20)

| Cluster size | T-value | x | y | z | Hemisphere | Region |
| --- | --- | --- | --- | --- | --- | --- |
| 686 | 10.38 | -28 | -82 | 34 | Left | Superior Occipital Gyrus / Lateral Occipital Cortex, superior / BA19 |
| 822 | 10.36 | 14 | -76 | -24 | Right | Cerebellum |
| 502 | 9.73 | 0 | -64 | 56 | Bilateral | Precuneus |
| 112 | 9.57 | 12 | -78 | 40 | Right | Precuneus |
| 538 | 9.42 | 10 | -34 | 48 | Right | Cingulate Gyrus, posterior division extended to precentral gyrus |
| 112 | 8.77 | -20 | -28 | 68 | Left | Precentral Gyrus |
| 161 | 8.66 | -44 | -38 | 54 | Left | Superior Parietal Lobule |
| 62 | 8.64 | 0 | -82 | 28 | Bilateral | Cuneal Cortex |
| 106 | 8.3 | 22 | -26 | 64 | Right | Precentral Gyrus |
| 419 | 8.25 | 30 | -54 | 50 | Right | Superior Parietal Lobule |
| 231 | 8.16 | 32 | -80 | 28 | Right | Superior Occipital Gyrus / Lateral Occipital Cortex, superior / BA19 |
| 65 | 6.55 | -8 | -78 | 36 | Left | Precuneus |
| 33 | 7.07 | 40 | -22 | 60 | Right | Precentral Gyrus |
| 29 | 6.55 | 56 | -48 | 36 | Right | Angular Gyrus |
| 25 | 6.6 | 10 | -76 | -12 | Right | Lingual Gyrus |
| 25 | 6.06 | 50 | -62 | 28 | Right | Lateral Occipital Cortex, superior / Middle Temporal Gyrus / Angular Gyrus |
| 24 | 7.01 | -50 | -14 | 50 | Left | Postcentral Gyrus |
| 24 | 6.31 | -42 | -78 | 26 | Left | Lateral Occipital Cortex, superior / Middle Occipital/Temporal Gyrus |
| 24 | 6.19 | 4 | -90 | -16 | Right | Cerebellum |
| 24 | 5.83 | -42 | -16 | 50 | Left | Precentral Gyrus |
| 28 | 5.62 | 0 | -54 | 6 | Midline | Cerebellum |

**Table S4.**
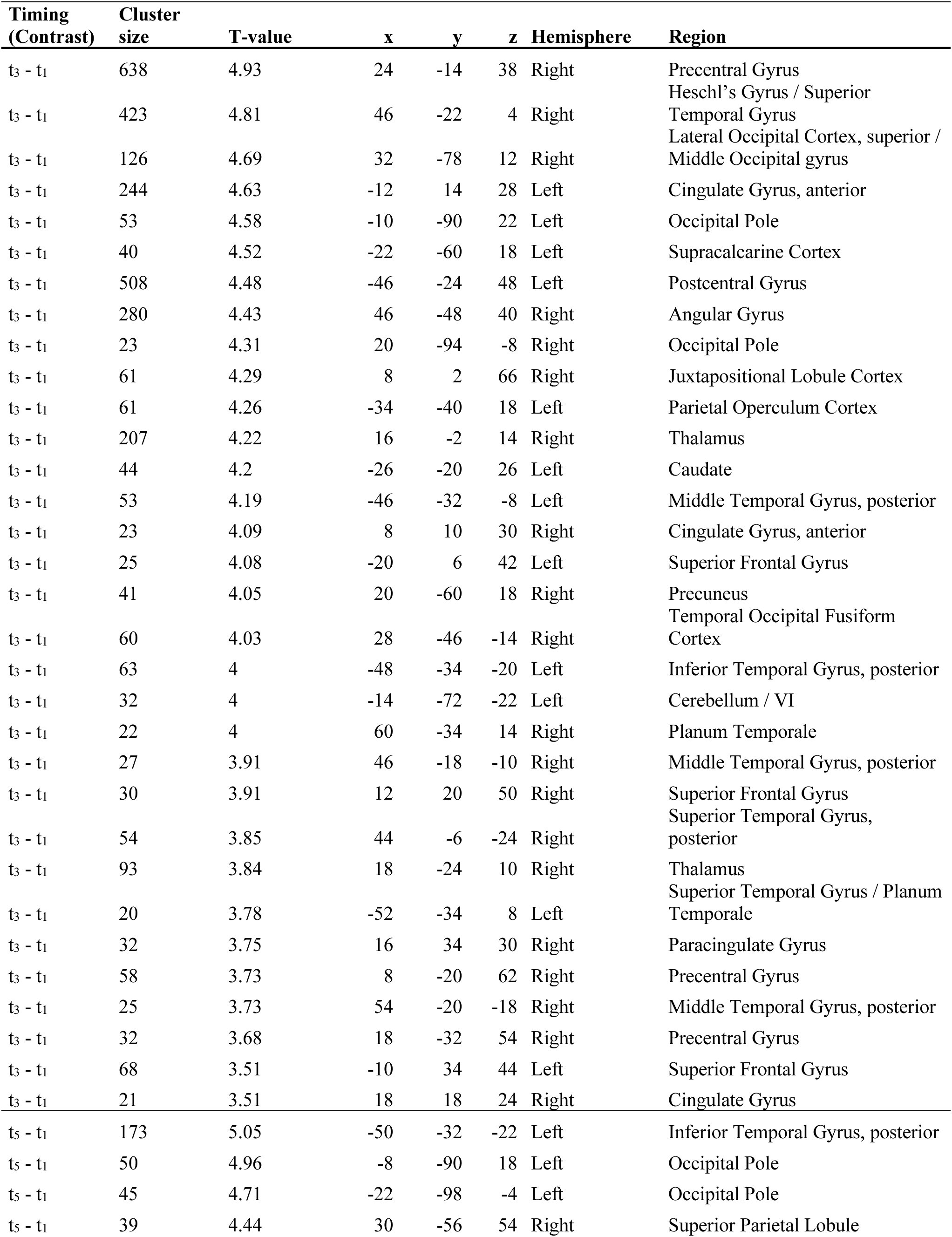
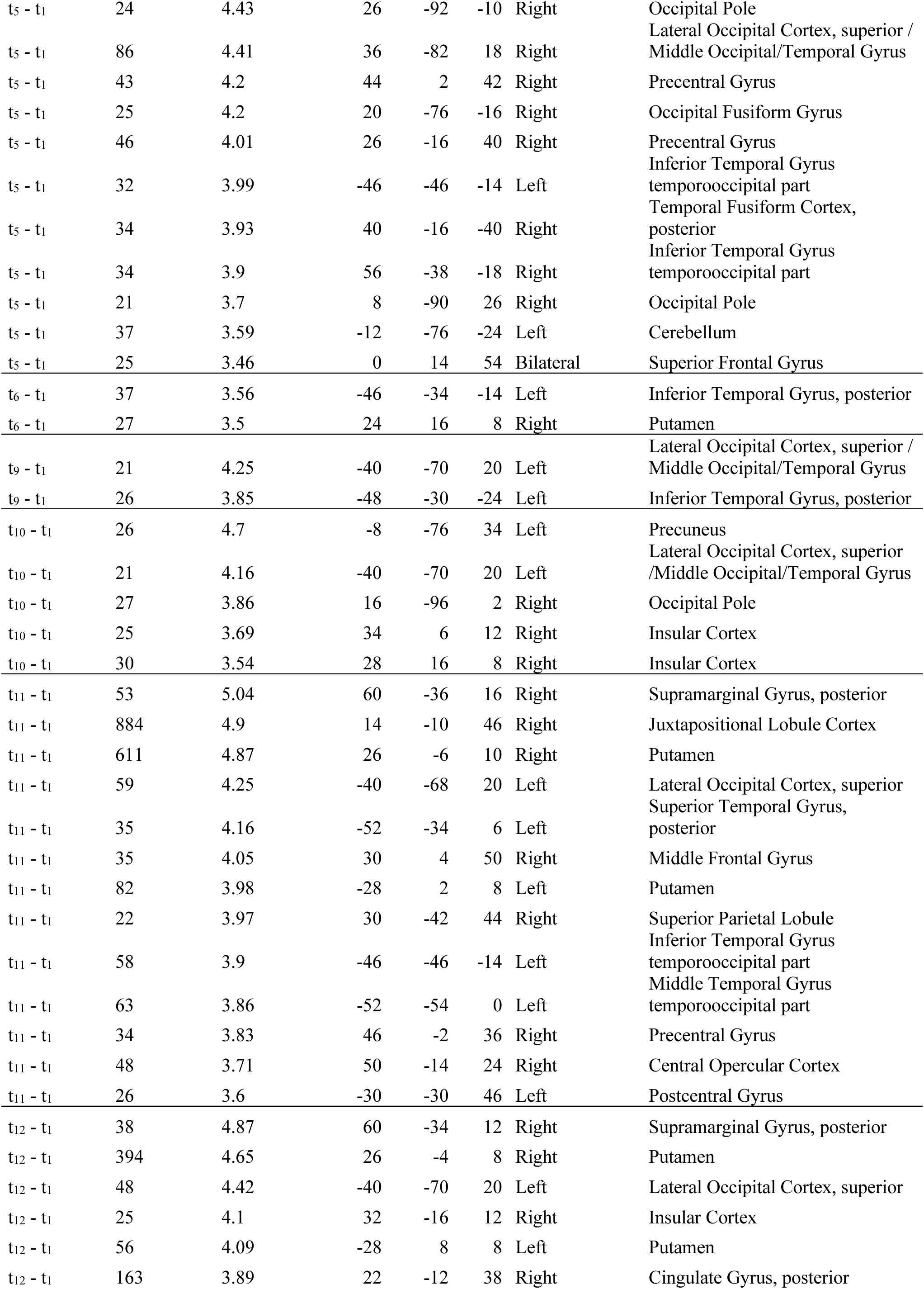
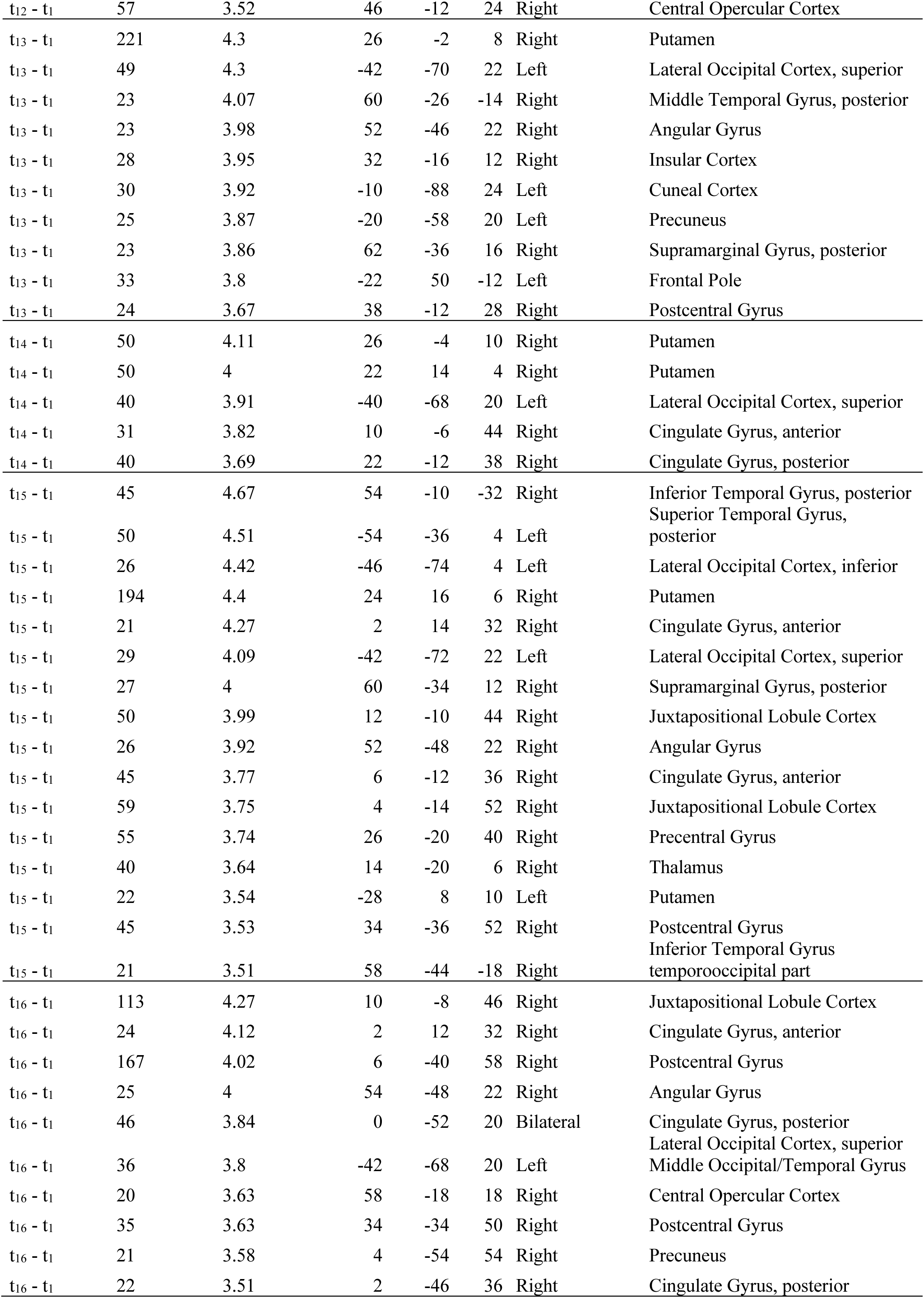
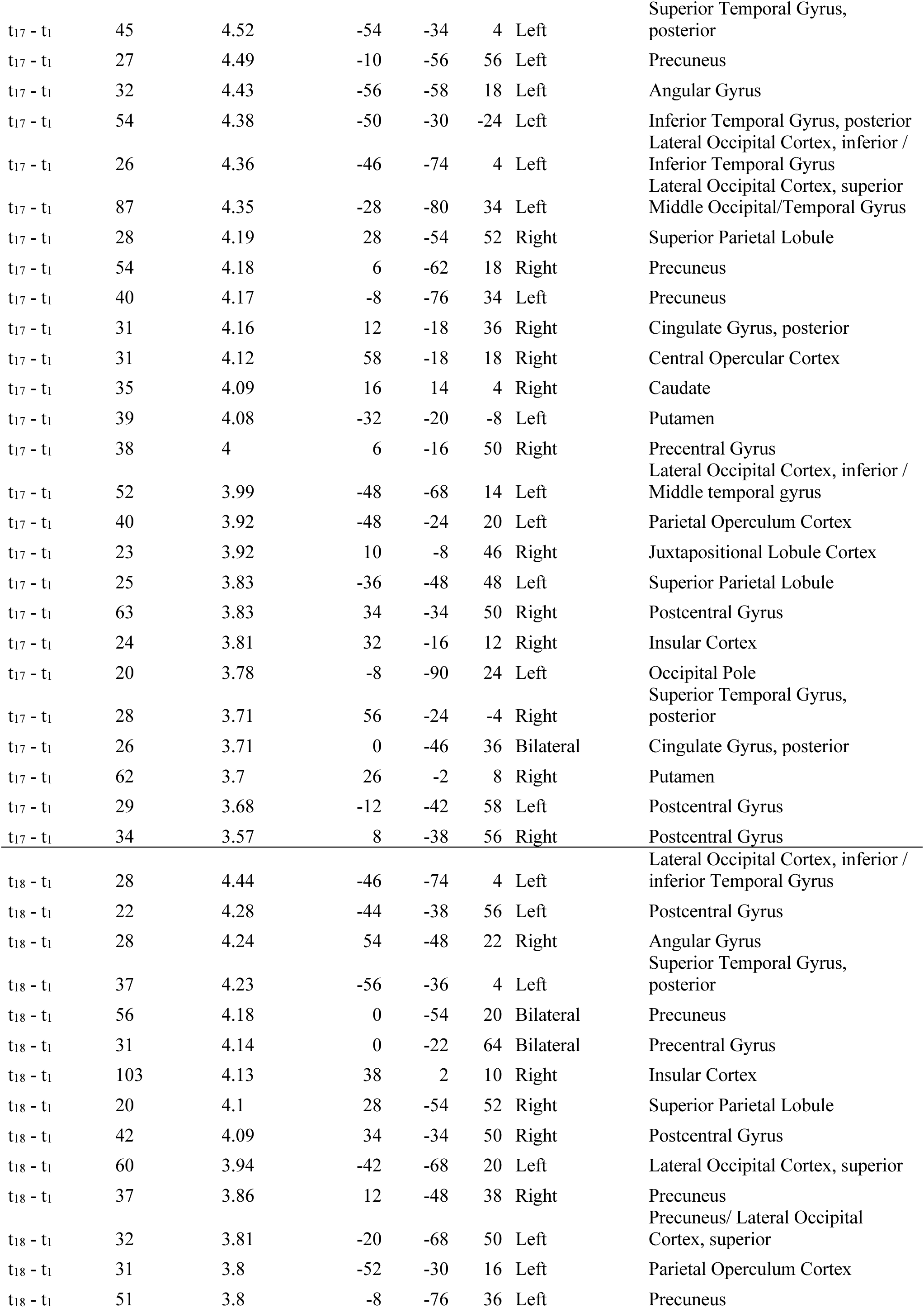
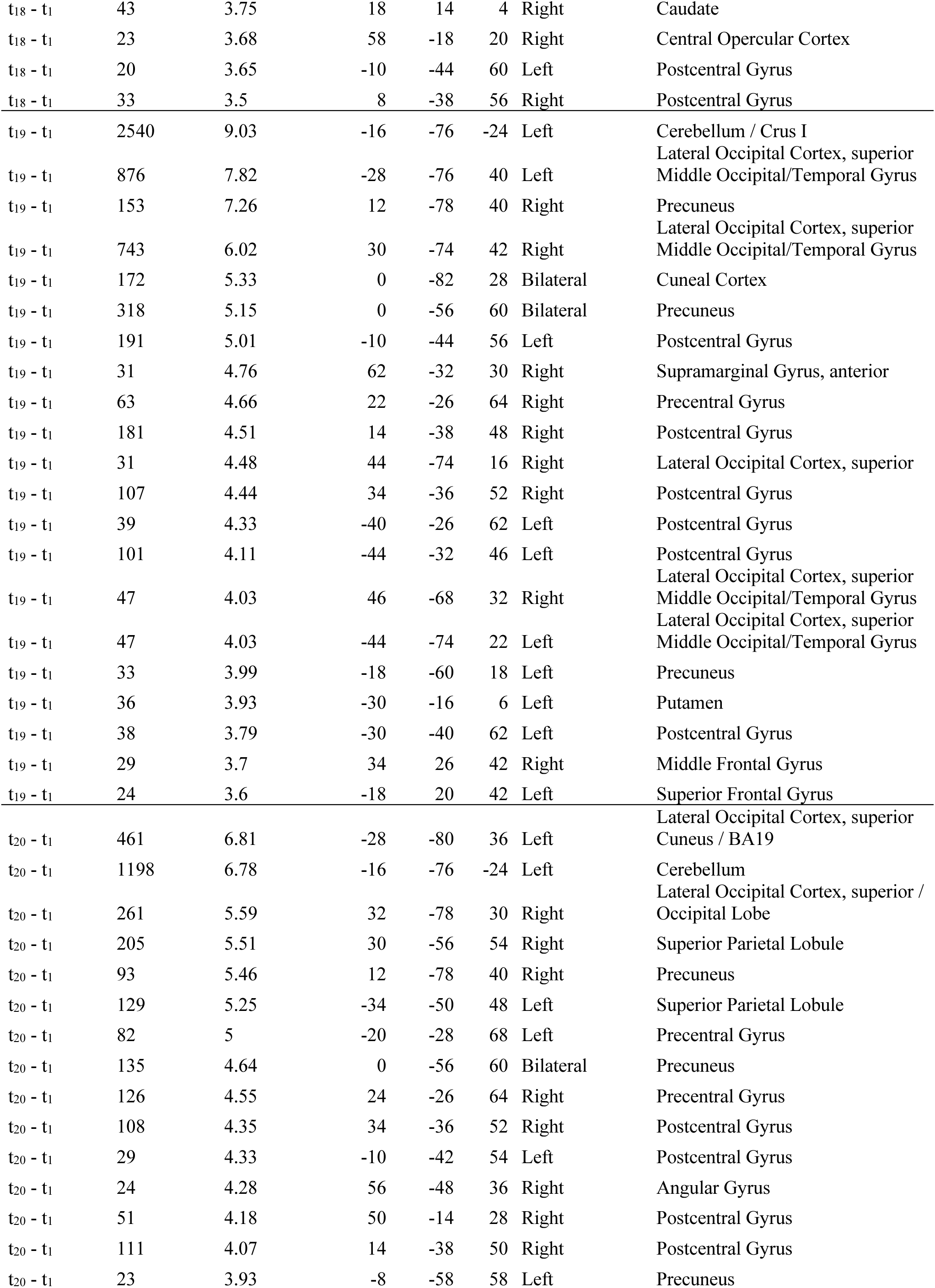
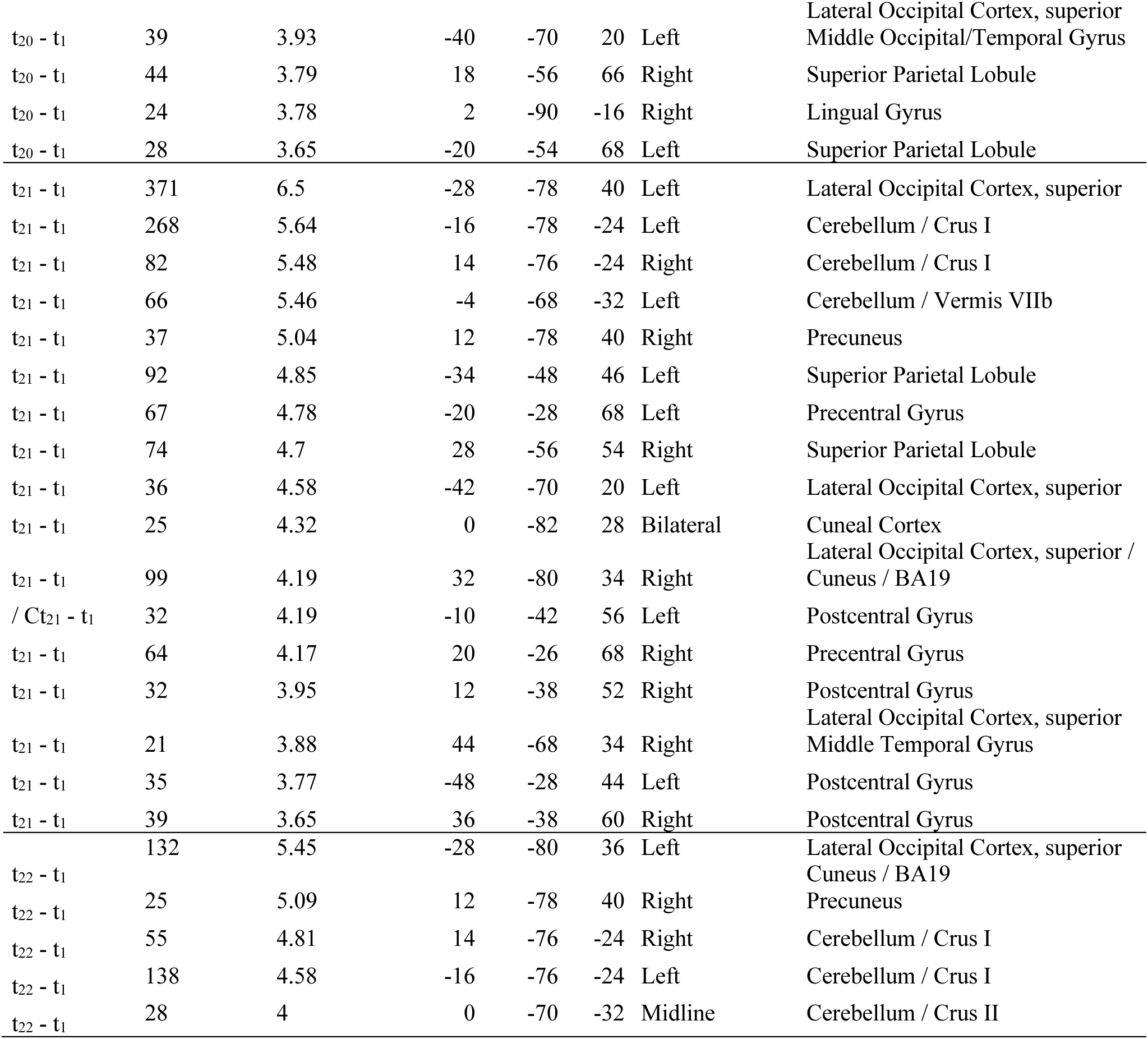
Temporal trajectories of learning-induced mean diffusivity (MD) changes by contrasting the MD change from baseline (t_1_) to each subsequent time point (t_2_-t_22_) against the corresponding change in the control condition. (*p*_uncorrected_<.001, k=20)

**Table S5.** Overlap statistics of left middle occipital/temporal gyrus across time.

| **time** | **Cluster size** | **Number of voxels in overlapping cluster** | **Percentage of the cluster** | **Percentage of the overlapping cluster** |
| --- | --- | --- | --- | --- |
| 9 | 21 | 15 | 71.42 | 88.23 |
| 10 | 21 | 12 | 57.14 | 70.58 |
| 11 | 59 | 16 | 27.11 | 94.11 |
| 12 | 48 | 17 | 35.41 | 100.0 |
| 13 | 49 | 17 | 34.69 | 100.0 |
| 14 | 40 | 14 | 35.0 | 82.35 |
| 15 | 29 | 16 | 55.17 | 94.11 |
| 16 | 35 | 14 | 40.0 | 82.35 |
| 17 | 52 | 16 | 30.76 | 94.11 |
| 18 | 60 | 16 | 26.67 | 94.11 |
| 19 | 47 | 12 | 25.53 | 70.58 |
| 20 | 39 | 12 | 30.76 | 70.58 |
| 21 | 36 | 14 | 38.89 | 82.35 |

**Table S6.** Post-hoc Pairwise Contrasts From Linear Mixed-Effects Models: Time Points Compared to Baseline (T1), Separately for Control and Learning Groups.

| Minute (time) | session | Contrast (Control) | Estimate | SE | df | *t* | *p* |
| --- | --- | --- | --- | --- | --- | --- | --- |
| -15.81 | base | t<sub>2</sub> - t<sub>1</sub> | $4.82 \times 10^{-07}$ | $1.37 \times 10^{-06}$ | 2095 | 0.353 | 1.0000 |
| 0 | task | t<sub>3</sub> - t<sub>1</sub> | $3.22 \times 10^{-07}$ | $1.70 \times 10^{-06}$ | 2095 | 0.189 | 1.0000 |
| 3.83 | task | t<sub>4</sub> - t<sub>1</sub> | $3.67 \times 10^{-06}$ | $1.70 \times 10^{-06}$ | 2095 | 2.153 | 0.6292 |
| 7.66 | task | t<sub>5</sub> - t<sub>1</sub> | $4.67 \times 10^{-06}$ | $1.70 \times 10^{-06}$ | 2095 | 2.742 | 0.1294 |
| 11.5 | task | t<sub>6</sub> - t<sub>1</sub> | $3.52 \times 10^{-06}$ | $1.70 \times 10^{-06}$ | 2095 | 2.068 | 0.7365 |
| 16.96 | task | t<sub>7</sub> - t<sub>1</sub> | $3.64 \times 10^{-07}$ | $1.74 \times 10^{-06}$ | 2095 | 0.209 | 1.0000 |
| 20.8 | task | t<sub>8</sub> - t<sub>1</sub> | $-1.12 \times 10^{-06}$ | $1.74 \times 10^{-06}$ | 2095 | -0.645 | 1.0000 |
| 24.63 | task | t<sub>9</sub> - t<sub>1</sub> | $-1.26 \times 10^{-06}$ | $1.74 \times 10^{-06}$ | 2095 | -0.726 | 1.0000 |
| 28.46 | task | t<sub>10</sub> - t<sub>1</sub> | $-1.87 \times 10^{-06}$ | $1.74 \times 10^{-06}$ | 2095 | -1.073 | 1.0000 |
| 32.3 | task | t<sub>11</sub> - t<sub>1</sub> | $1.22 \times 10^{-06}$ | $1.70 \times 10^{-06}$ | 2095 | 0.718 | 1.0000 |
| 36.13 | task | t<sub>12</sub> - t<sub>1</sub> | $1.08 \times 10^{-06}$ | $1.70 \times 10^{-06}$ | 2095 | 0.631 | 1.0000 |
| 39.96 | task | t<sub>13</sub> - t<sub>1</sub> | $1.65 \times 10^{-06}$ | $1.70 \times 10^{-06}$ | 2095 | 0.971 | 1.0000 |
| 43.8 | task | t<sub>14</sub> - t<sub>1</sub> | $3.35 \times 10^{-06}$ | $1.70 \times 10^{-06}$ | 2095 | 1.964 | 0.8933 |
| 49.26 | task | t<sub>15</sub> - t<sub>1</sub> | $-2.49 \times 10^{-06}$ | $1.74 \times 10^{-06}$ | 2095 | -1.430 | 1.0000 |
| 53.1 | task | t<sub>16</sub> - t<sub>1</sub> | $-1.29 \times 10^{-06}$ | $1.74 \times 10^{-06}$ | 2095 | -0.743 | 1.0000 |
| 56.933 | task | t<sub>17</sub> - t<sub>1</sub> | $5.10 \times 10^{-07}$ | $1.74 \times 10^{-06}$ | 2095 | 0.294 | 1.0000 |
| 60.766 | task | t<sub>18</sub> - t<sub>1</sub> | $1.83 \times 10^{-07}$ | $1.74 \times 10^{-06}$ | 2095 | 0.106 | 1.0000 |
| 116.13 | post | t<sub>35</sub> - t<sub>1</sub> | $1.44 \times 10^{-06}$ | $1.37 \times 10^{-06}$ | 2095 | 1.054 | 1.0000 |
| 119.96 | post | t<sub>36</sub> - t<sub>1</sub> | $6.41 \times 10^{-07}$ | $1.37 \times 10^{-06}$ | 2095 | 0.469 | 1.0000 |
| 123.8 | post | t<sub>37</sub> - t<sub>1</sub> | $3.61 \times 10^{-07}$ | $1.37 \times 10^{-06}$ | 2095 | 0.264 | 1.0000 |
| 127.63 | post | t<sub>38</sub> - t<sub>1</sub> | $-4.63 \times 10^{-07}$ | $1.37 \times 10^{-06}$ | 2095 | -0.339 | 1.0000 |

| Minute (time) |  | Contrast<br>(Learning) | Estimate | SE | df | *t* | *p* |
| --- | --- | --- | --- | --- | --- | --- | --- |
| -15.8 | base | $t_2 - t_1$ | $-1.15 \times 10^{-06}$ | $9.67 \times 10^{-07}$ | 2095 | -1.190 | 0.2342 |
| 0 | task | $t_3 - t_1$ | $-2.71 \times 10^{-06}$ | $1.22 \times 10^{-06}$ | 2095 | -2.228 | 0.0888 |
| 3.83 | task | $t_4 - t_1$ | $-2.85 \times 10^{-06}$ | $1.22 \times 10^{-06}$ | 2095 | -2.342 | 0.0888 |
| 7.666 | task | $t_5 - t_1$ | $-4.67 \times 10^{-06}$ | $1.22 \times 10^{-06}$ | 2095 | -3.837 | 0.0022 |
| 11.5 | task | $t_6 - t_1$ | $-3.41 \times 10^{-06}$ | $1.22 \times 10^{-06}$ | 2095 | -2.799 | 0.0463 |
| 16.96 | task | $t_7 - t_1$ | $-2.53 \times 10^{-06}$ | $1.22 \times 10^{-06}$ | 2095 | -2.081 | 0.0888 |
| 20.8 | task | $t_8 - t_1$ | $-4.33 \times 10^{-06}$ | $1.22 \times 10^{-06}$ | 2095 | -3.559 | 0.0049 |
| 24.63 | task | $t_9 - t_1$ | $-3.52 \times 10^{-06}$ | $1.22 \times 10^{-06}$ | 2095 | -2.893 | 0.0386 |
| 28.46 | task | $t_{10} - t_1$ | $-3.41 \times 10^{-06}$ | $1.22 \times 10^{-06}$ | 2095 | -2.801 | 0.0463 |
| 32.3 | task | $t_{11} - t_1$ | $-2.97 \times 10^{-06}$ | $1.22 \times 10^{-06}$ | 2095 | -2.441 | 0.0883 |
| 36.13 | task | $t_{12} - t_1$ | $-5.06 \times 10^{-06}$ | $1.22 \times 10^{-06}$ | 2095 | -4.156 | 0.0007 |
| 39.96 | task | $t_{13} - t_1$ | $-6.08 \times 10^{-06}$ | $1.22 \times 10^{-06}$ | 2095 | -4.998 | <.0001 |
| 43.8 | task | $t_{14} - t_1$ | $-5.06 \times 10^{-06}$ | $1.22 \times 10^{-06}$ | 2095 | -4.156 | 0.0007 |
| 49.26 | task | $t_{15} - t_1$ | $-2.89 \times 10^{-06}$ | $1.22 \times 10^{-06}$ | 2095 | -2.372 | 0.0888 |
| 53.1 | task | $t_{16} - t_1$ | $-5.49 \times 10^{-06}$ | $1.22 \times 10^{-06}$ | 2095 | -4.514 | 0.0002 |
| 56.93 | task | $t_{17} - t_1$ | $-5.46 \times 10^{-06}$ | $1.22 \times 10^{-06}$ | 2095 | -4.490 | 0.0002 |
| 60.76 | task | $t_{18} - t_1$ | $-4.63 \times 10^{-06}$ | $1.22 \times 10^{-06}$ | 2095 | -3.805 | 0.0023 |
| 74.6 | post | $t_{19} - t_1$ | $-5.36 \times 10^{-06}$ | $1.22 \times 10^{-06}$ | 2095 | -4.404 | 0.0003 |
| 78.43 | post | $t_{20} - t_1$ | $-5.11 \times 10^{-06}$ | $1.22 \times 10^{-06}$ | 2095 | -4.200 | 0.0006 |
| 80.48 | post | $t_{21} - t_1$ | $-4.80 \times 10^{-06}$ | $1.22 \times 10^{-06}$ | 2095 | -3.945 | 0.0015 |
| 82.26 | post | $t_{22} - t_1$ | $-7.28 \times 10^{-06}$ | $1.22 \times 10^{-06}$ | 2095 | -5.980 | <.0001 |
| 84.31 | post | $t_{23} - t_1$ | $-6.12 \times 10^{-06}$ | $1.22 \times 10^{-06}$ | 2095 | -5.026 | <.0001 |
| 86.1 | post | $t_{24} - t_1$ | $-5.36 \times 10^{-06}$ | $1.22 \times 10^{-06}$ | 2095 | -4.403 | 0.0003 |
| 88.15 | post | $t_{25} - t_1$ | $-3.36 \times 10^{-06}$ | $1.22 \times 10^{-06}$ | 2095 | -2.761 | 0.0463 |
| 91.98 | post | $t_{26} - t_1$ | $-4.14 \times 10^{-06}$ | $1.22 \times 10^{-06}$ | 2095 | -3.406 | 0.0080 |
| 95.98 | post | $t_{27} - t_1$ | $-4.86 \times 10^{-06}$ | $1.22 \times 10^{-06}$ | 2095 | -3.997 | 0.0013 |
| 99.65 | post | $t_{28} - t_1$ | $-4.82 \times 10^{-06}$ | $1.22 \times 10^{-06}$ | 2095 | -3.963 | 0.0015 |
| 100.8 | post | $t_{29} - t_1$ | $-5.62 \times 10^{-06}$ | $1.22 \times 10^{-06}$ | 2095 | -4.619 | 0.0001 |
| 103.48 | post | $t_{30} - t_1$ | $-4.63 \times 10^{-06}$ | $1.22 \times 10^{-06}$ | 2095 | -3.805 | 0.0023 |
| 104.63 | post | $t_{31} - t_1$ | $-4.40 \times 10^{-06}$ | $1.22 \times 10^{-06}$ | 2095 | -3.620 | 0.0042 |
| 107.31 | post | $t_{32} - t_1$ | $-3.63 \times 10^{-06}$ | $1.22 \times 10^{-06}$ | 2095 | -2.980 | 0.0321 |
| 108.11 | post | $t_{33} - t_1$ | $-5.30 \times 10^{-06}$ | $1.22 \times 10^{-06}$ | 2095 | -4.359 | 0.0003 |
| 112.3 | post | $t_{34} - t_1$ | $-6.53 \times 10^{-06}$ | $1.22 \times 10^{-06}$ | 2095 | -5.369 | <.0001 |
| 116.13 | post | $t_{35} - t_1$ | $-5.12 \times 10^{-06}$ | $9.67 \times 10^{-07}$ | 2095 | -5.298 | <.0001 |
| 119.96 | post | $t_{36} - t_1$ | $-5.73 \times 10^{-06}$ | $9.67 \times 10^{-07}$ | 2095 | -5.922 | <.0001 |
| 123.8 | post | $t_{37} - t_1$ | $-6.82 \times 10^{-06}$ | $9.67 \times 10^{-07}$ | 2095 | -7.050 | <.0001 |
| 127.63 | post | $t_{38} - t_1$ | $-5.09 \times 10^{-06}$ | $9.67 \times 10^{-07}$ | 2095 | -5.264 | <.0001 |

*Note.* SE = Standard Error; df = degrees of freedom; *p*-values are adjusted using the Holm method.

